# A Teensy microcontroller-based interface for optical imaging camera control during behavioral experiments

**DOI:** 10.1101/475350

**Authors:** Michael Romano, Mark Bucklin, Dev Mehrotra, Robb Kessel, Howard Gritton, Xue Han

**Author notes:** **Corresponding author**. Please send to Xue Han. 44 Cummington Street, Boston, MA 02215.

## Abstract

**Background:** Systems neuroscience experiments often require the integration of precisely timed data acquisition and behavioral monitoring. While specialized commercial systems have been designed to meet various needs of data acquisition and device control, they often fail to offer flexibility to interface with new instruments and variable behavioral experimental designs.

**New method:** We developed a Teensy 3.2 microcontroller-based interface that offers high-speed, precisely timed behavioral data acquisition and digital and analog outputs for controlling sCMOS cameras and other devices.

**Results:** We demonstrate the flexibility and the temporal precision of the Teensy interface in two experimental settings. We first used the Teensy interface for reliable recordings of an animal’s directional movement on a spherical treadmill, while delivering repeated digital pulses that can be used to control image acquisition from a sCMOS camera. In another example, we used the Teensy interface to control temporally precise delivery of an auditory stimulus and a gentle eye puff in a trace conditioning eye blink behavioral paradigm, while delivering repeated digital pulses to initiate camera image acquisition.

**Comparison with existing methods:** This interface allows high-speed and temporally precise digital data acquisition and device control during diverse behavioral experiments.

**Conclusion:** The Teensy interface, consisting of a Teensy 3.2 and custom software functions, provides a temporally precise, low-cost, and flexible platform to integrate sCMOS camera control into behavioral experiments.

## 1. Introduction

Recent advances in sCMOS camera technology and genetically encoded neural activity indicators enable large scale fluorescence imaging of thousands of individual cells’ activity during behavior (Mohammed, et al. 2016, Nguyen, et al. 2016). One key technical aspect of neural network analysis during behavior is temporal precision, where neural activities need to be precisely aligned with behavioral features. However, it has been difficult to easily integrate sCMOS cameras, deployed in large scale calcium imaging studies, with devices needed to monitor and control behavioral experiments. Traditional Analog/Digital interfaces are often operated by programs, such as MATLAB, that offer a wide range of applications. However, using MATLAB or other PC-based programs can lead to undesired temporal delays, as the PC operating system needs to balance the demands of many system operations at once.

Over the last decade, microcontrollers traditionally marketed to hobbyists have gained popularity across a variety of scientific fields (Chen and Li 2017, D’Ausilio 2012, Husain, Hadad and Zainal Alam 2016, Sanders and Kepecs 2014). For example, Arduino microcontrollers have recently been integrated into two-photon imaging experiments (Micallef, et al. 2017, Takahashi, et al. 2016, Wilms and Häusser 2015). Microcontrollers are small, low-cost, and capable of delivering digital outputs with microsecond time precision. Arduino, which utilizes user-friendly, open-source software functions, was the first major microcontroller to gain substantial popularity. Recently, Teensy 3.2 microcontrollers were developed, which have all the key features of the current version of the standard Arduino microcontroller (Arduino Uno Rev3), as well as the additional feature of delivering analog output. Teensy microcontrollers utilize the same open-source Arduino software environment, and remain easy to program (D’Ausilio 2012). Because of the simplicity of microcontrollers and their temporal precisions, microcontrollers represent an attractive solution to precisely record data and monitor experimental progress.

Here, we demonstrate and characterize a flexible Teensy 3.2-based interface for temporally precise data acquisition and delivery of analog and digital signals, during a voluntary movement tracking experiment and trace conditioning eye blink learning experiment. The Teensy interface can deliver digital pulses with microsecond precision to initiate individual image frame capture using the camera’s external trigger settings at a desired frequency, while simultaneously collecting animal behavioral data. We also demonstrate the ability of the Teensy interface to generate analog sound waveforms to drive a speaker for a trace conditioning eye blink experiment. Together, these results demonstrate that the Teensy interface, consisting of a Teensy microcontroller and a set of custom software functions, offers a flexible, accurate, and user-friendly environment for imaging experiments during behavior.

## 2. Methods

### 2.1 Construction of Teensy 3.2 boards

The two experimental designs are shown in Figure 1. The specialty components required to build these designs are shown in Tables 1 and 2. In both experiments, a Teensy 3.2 (PJRC.COM, LLC, part #: TEENSY32) (Figure 1A), or a Teensy 3.2 soldered to a prop shield (PJRC.COM, part #: PROP_SHIELD) (Figure 1B), is mounted on top of a standard printed circuit board (PCB) (for example: Digi-Key, part #: V2010-ND) via standard female headers (such as SparkFun Electronics, PRT-00115). Female headers were then soldered to the PCB for stability. Output from the Teensy was directed from pins on the female headers to standard SMA connectors (such as: Digi-Key, part # CON-SMA-EDGE-S-ND) via 22 gauge wires (for example: Digi-Key, part #1528-1743-ND). Coaxial cables were then attached to the SMA connectors on the PCB to connect the Teensy to external devices. The Teensy was connected to a computer via a standard USB-microUSB cable (for example: Digi-Key, part # AE11229-ND). To easily upload code to the Teensy, we used PlatformIO (https://platformio.org/), an add-on to the widely-used Atom text editor (https://atom.io/), instead of the default Arduino programming environment. Code for each of the two experimental settings were uploaded separately to the Teensy prior to each experiment. To turn digital pins on and off, and also to change their modes to either “input” or “output”, we used a slightly modified version of the DigitalIO library provided by PlatformIO (version 1.0.0; currently maintained at: https://github.com/greiman/DigitalIO), which decreases the amount of time, which decreases the amount of time/ spent performing each of these actions. To easily set experiment-specific parameters for the Teensy, such as sampling frequencies, trial numbers and trial length, and the length of an experiment, we developed two simple MATLAB graphical user interfaces, one for each experiment.

**Table 1.**
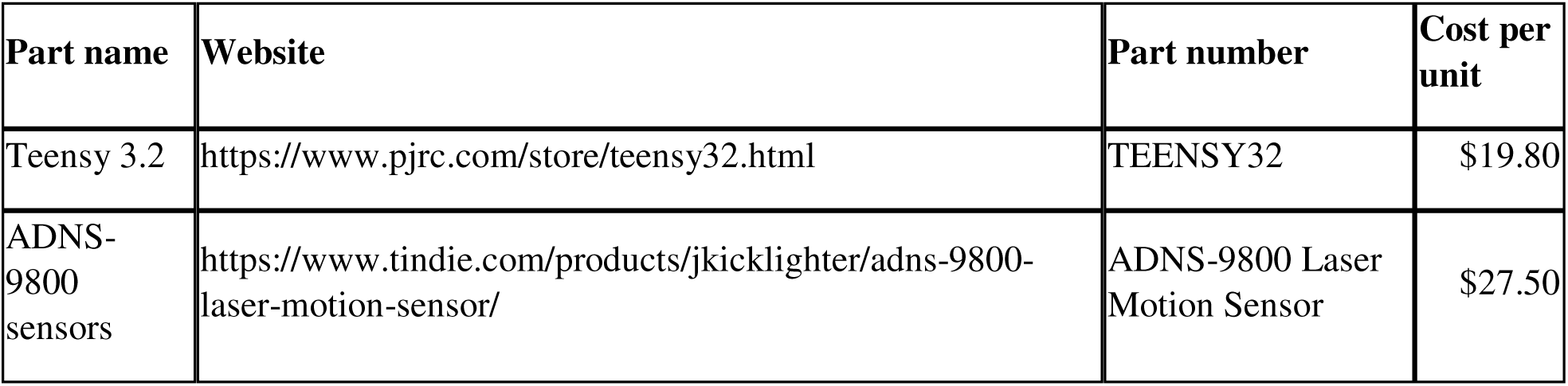
Specialty components necessary to build a motor output system

| Part name | Website | Part number | Cost per unit |
| --- | --- | --- | --- |
| Teensy 3.2 | https://www.pjrc.com/store/teensy32.html | TEENSY32 | \$19.80 |
| ADNS-9800 sensors | https://www.tindie.com/products/jkicklighter/adns-9800-laser-motion-sensor/ | ADNS-9800 Laser Motion Sensor | \$27.50 |

**Table 2.** Specialty components necessary to build a tone-puff system.

| Part name | Website | Part number | Cost per unit |
| --- | --- | --- | --- |
| Teensy 3.2 | https://www.pjrc.com/store/teensy32.html | TEENSY32 | \$19.80 |
| 14x1 Double insulator pins | https://www.pjrc.com/store/header_14x1_d.html | HEADER_14x1_D | \$0.85 |
| Prop shield | https://www.pjrc.com/store/prop_shield.html | PROP_SHIELD | \$19.50 |

**Figure 1.**
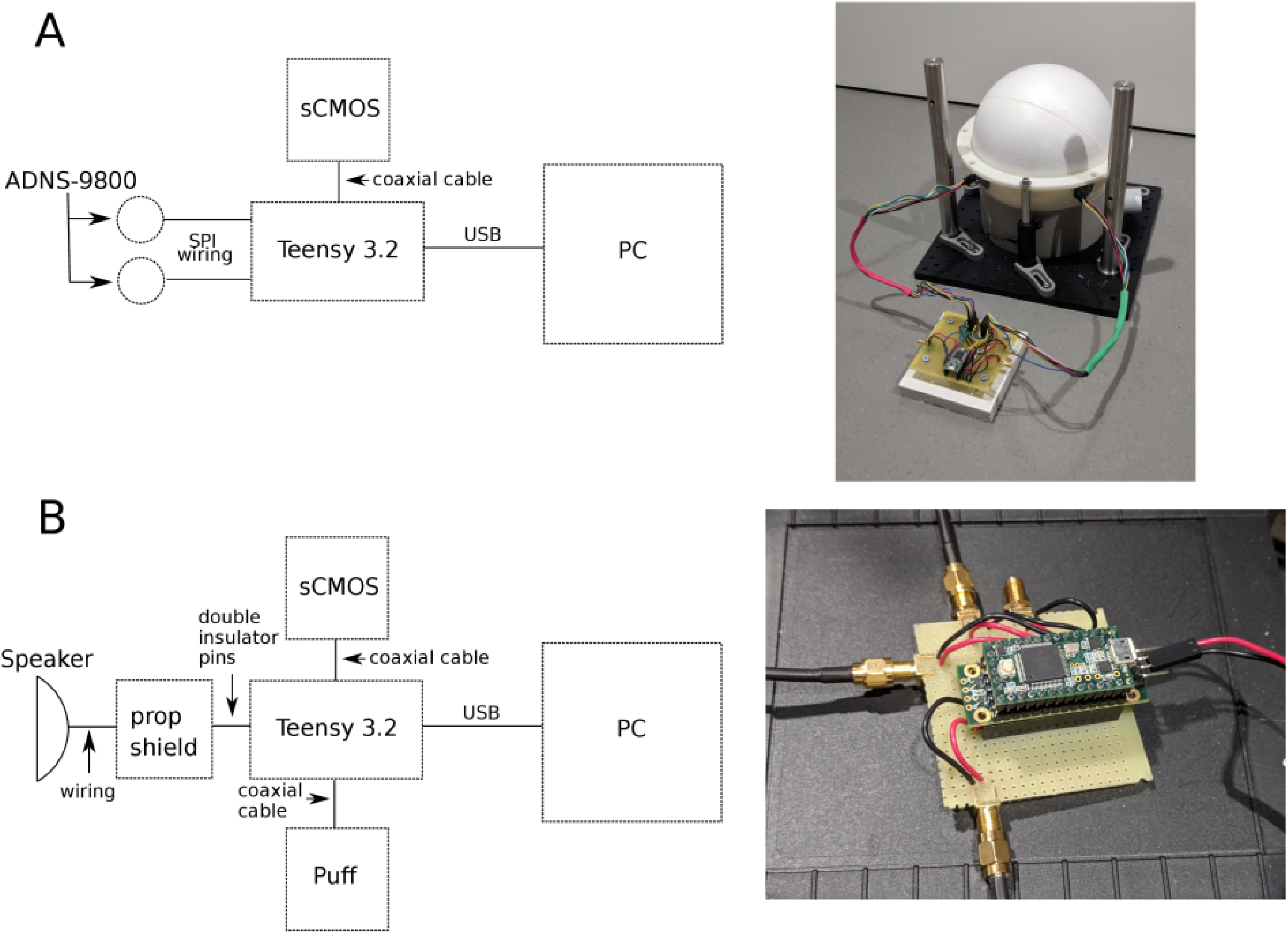
Diagrams of the two experimental device arrangements using a Teensy interface. **A** Motion tracking experiment design. This design consists of a Teensy 3.2 connected to two ADNS-9800 sensors via serial-peripheral interfaces, and a sCMOS camera through a coaxial cable. Every 50 milliseconds, a digital pulse was sent to initiate an image frame capture from a sCMOS camera. Simultaneously, the Teensy interface acquired motion data from both ADNS sensors and sent them to a PC via a USB. **B** Trace eye blink conditioning experiment design. This design consists of consists of a Teensy 3.2 connected to a speaker through a prop-shield that contains an amplifier. Every 50 milliseconds, a digital pulse was sent to initiate an image frame capture from a sCMOS camera. Simultaneously, the Teensy interface generated digital pulses to generate air puff and updated the status of the analog output to generate audio signals, and sent the timing of these signals to a PC via a USB.

### 2.2 Motion tracking experiment

In this experiment, we performed motion tracking using two ADNS-9800 gaming sensors (https://www.tindie.com/products/jkicklighter/adns-9800-laser-motion-sensor/, Tindie, part: <u>“NS-9800</u> <u>Laser Motion Sensor</u>”, see Table 1), while delivering digital pulses that can be used to trigger a sCMOS camera for image capture every 50 ms. The overall design of this experiment is shown in Figure 1A. A mouse was positioned on top of a buoyant Styrofoam ball floated by house air as described previously (Dombeck, et al. 2007). Two ADNS-9800 gaming sensors were positioned at the equator of the Styrofoam ball, at an angle of approximately 75 degrees from one another. For the counts per inch setting of the sensor, which determines the sensitivity of the sensors to external movement, we used a value of 3400 counts per inch.

The total distance travelled by the mouse at any time point was computed using the following equation:

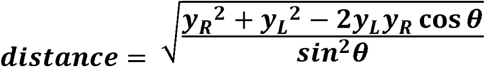

Where y_R_ and y_L_ are the y readings from the left and right sensors, and ***θ*** is the angle between the two sensors (75 degrees). We also acquired readings in the “x” direction from both sensors, which can be used to calculate rotation. Velocity was computed as the distance divided by the time between two adjacent readings, as measured by the Teensy. These two sensors were connected to the Teensy via simple serial peripheral interface (SPI) connections with insulated 22 gauge wires as shown in Figure 2A.

**Figure 2.**
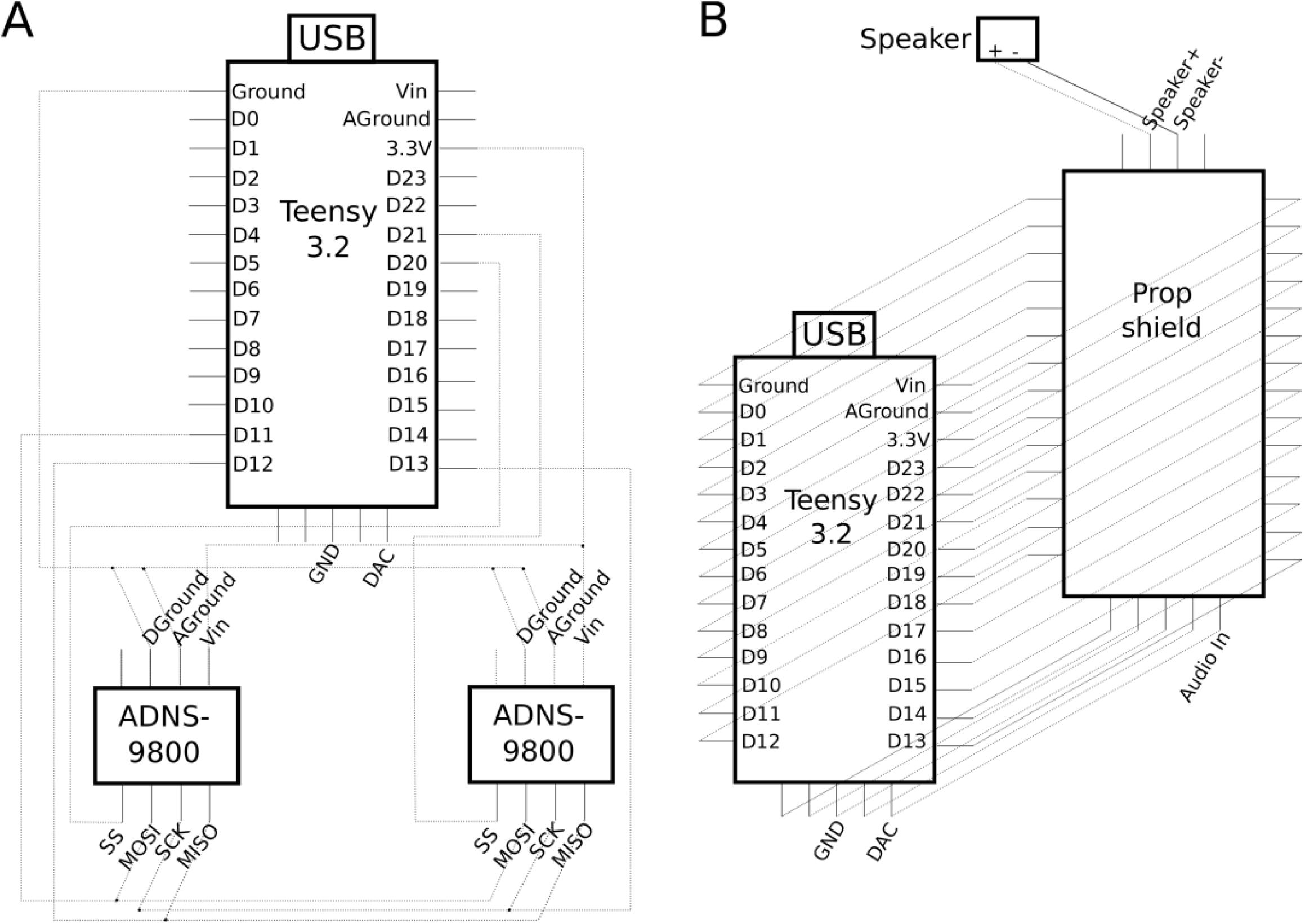
Electrical wiring schematics for the motion tracking experiment and the trace conditioning eye blink experiment **A.** The schematic of the wiring of a Teensy 3.2 to two ADNS-9800 sensors via serial peripheral interface connections (SPIs). Solid dots at intersections between dotted lines indicate electrical connections. Unused pins on the Teensy were not included in this schematic. The Teensy’s ground pin was connected to both AGround and DGround pins (analog and digital ground) on both ADNS-9800 sensors. The D11 pin (D = digital) was connected to both MOSI (“Master-Out, Slave-In”) pins, the D12 pin was connected to both MISO pins (“Master-In, Slave-Out”), the D13 pin was connected to both SCK pins (SPI Clock), and the 3.3V pin was connected to both Vin (voltage in) pins on the ADNS-9800 sensors. Finally, pins D20 and D21 were connected individually to each SS pin (Slave Select) on the ADNS-9800 sensors. The DAC pin (digital to analog converter or the analog output pin) is also shown. **B** The schematic of the wiring of a Teensy 3.2, a prop shield, and an external speaker. Dotted lines indicate connections. Connections between the Teensy and prop shield were made using 14×1 double insulated pins according to the manufacturer’s instruction (https://www.pjrc.com/store/prop_shield.html), and the prop shield audio output was connected to the speaker using 22 gauge wire. We highlight that that the Teensy DAC pin is connected to the “Audio In” pin on the prop shield, both of which are labeled. Additional pins utilized by the prop shield for amplification were not labeled.

To control experimental timing, we utilized the “IntervalTimer” function, unique to the standard Teensy library, which can repeatedly call a function at specified intervals. We set the “IntervalTimer” to be 50,000 microseconds (50 ms) or 20 Hz in our experiment. Using IntervalTimer, we repeatedly called a function that sent the accumulated displacement of the motion sensor readings to the attached PC. We acquired the x and y displacement readings from each sensor with freely available functions on GitHub (https://github.com/markbucklin/NavigationSensor), which read accumulated displacement from the “motion burst” register of each sensor. After reading the motion sensor, a digital “on” pulse that lasted for 1 ms was sent out of a digital pin designed to initiate an image frame capture from a sCMOS camera. To characterize the temporal precision of different digital pulses generated by the custom script using the “IntervalTimer” function, we recorded the digital outputs with a commercial system (Tucker Davis Technologies RZ5D (TDT RZ5D)) at 3051.76 Hz.

### 2.3 Trace eye blink conditioning experiment

In this experiment, a Teensy was programmed to deliver outputs capable of eliciting a sound and initiating an eye puff, while delivering digital pulses that can be used to trigger a sCMOS camera for image capture every 50 ms. The overall design of this experiment is shown in Figure 1B. To deliver an audible sound through the Teensy, we used a Teensy prop shield module (PJRC.COM, LLC., part #: PROP_SHIELD) to amplify analog output (shown in Figure 2B as pin A14). This add-on component can drive speakers with resistances up to 8 ohms. The prop shield was soldered to the bottom of the Teensy with 14×1 double insulator pins (PJRC.COM, LLC., part #: HEADER_14×1_D), and the output was connected to a speaker, as shown in Figure 2B. The Teensy was then mounted onto the female headers separated by the prop shield, as shown in Figure 1B. The camera and air valve for the eye puff were attached to the microcontroller through coaxial cables (Figures 1B and 2B), and the speaker was connected with 22 gauge wire to the prop shield.

We used the Teensy Audio library function “AudioSynthWaveformSine” to generate tones. This function continuously outputs a sine wave with a sampling rate of 44.1 kHz from the analog pin. We first initialized the tone, in this case a 9500 Hz sine wave, at the beginning of each experiment, but set the amplitude to “0”, so that the tone was off. At the desired time, we switched the amplitude to 0.05 (out of a maximum of 1) to generate an audible tone. The value of 0.05 generated a tone of approximately 75 dB with our amplifier and speaker settings.

We used the “elapsedMicros” function to control the timing of the experiment. elapsedMicros offers precise timing like “IntervalTimer”, and additionally allows for simultaneous use of the Audio library. This experiment is trial-based, and each trial consisted of an 11.1 second long baseline period, a 700ms long tone, a 250ms long delay period, a 100ms long puff period, and a 7.85 second long post-puff period. Using an “elapsedMicros” timer, we repeatedly called a function that updated the status of each digital and analog output every 50 ms based on the trial structure of the task, and then turned on the digital output directed to the sCMOS camera for 1ms every 50ms.

To characterize the temporal precision of different digital pulses generated by the custom scripts, we recorded the digital outputs with a commercial system (Tucker Davis Technologies RZ5D (TDT RZ5D)) at 3051.76 Hz, and the analog sine wave output at 24414.0625 Hz without additional amplification. To determine the onset of the analog audio signal, the unamplified analog output from the Teensy was first high-pass filtered at 1 kHz using a 6^th^-order, zero-phase Butterworth digital filter (MATLAB command “filtfilt”). We then estimated the instantaneous amplitude of the 9500 Hz sine wave at each time point using the Hilbert transform of the filtered signal. The first time point where the amplitude rose above 0.005 was considered the onset of the analog signal, and the subsequent time point where it dropped below 0.005 was considered the offset. To compare the onset of the analog signal to the timing of digital pulses, we utilized the continuous voltage output from the digital pin for consistency. To acquire the digital pulse onset from the continuous signal, we thresholded this continuous voltage output at a value of 1 V, and took the first time point where the continuous voltage exceeded 1 V to be the digital pulse onset.

### 2.4 Statistics

Statistics were performed in MATLAB. Linear models were constructed using the “fitlm” function in MATLAB 2017b, using a Lenovo ThinkPad T450 with 16 GB of RAM..

### 2.5 Code availability

All code is located at GitHub (https://github.com/mfromano/micro-control), which will be made public upon publication.

## 3. Results

Microcontrollers such as Arduino microcontrollers have gained popularity in neuroscience research due to their user-friendly interface, open-source software environment, and their flexibility of device integration (Chen and Li 2017, D’Ausilio 2012, Micallef, et al. 2017). Recently, the Teensy 3.2 has been developed, which has an analog output, a major improvement over the popular Arduino Uno. Teensy devices also have a comprehensive Audio library, as well as the “IntervalTimer” and the “elapsedMicros” functions capable of generating precisely timed events repeatedly. Here, we present a Teensy-based interface to integrate frame-by-frame image capture with behavioral experimental control and data acquisition.

### 3.1 Motion tracking experiment

In this experiment (Figure 3A), we recorded a mouse running on a spherical treadmill for 10 minutes. Motion data was acquired at 20 Hz concomitantly with digital outputs that can be used to trigger individual image frame capture from a sCMOS camera. To measure locomotion from awake head fixed mice, we used the Teensy interface to record from two ADNS-9800 motion sensors (Figures 1A and 2A). ADNS-9800 sensor boards are low cost, and can measure up to 8200 counts per inch, allowing for sensitive measurement of mouse movement relative to other tracking devices. For example, standard computer mice, such as the Logitech M100 (Logitech, PN: 910-001601), measure up to 1000 counts per inch, making the ADNS-9800 sensor over 8 times more precise. For these experiments we affixed ADNS- 9800 sensors to the spherical treadmill and wired them to the Teensy as demonstrated in Figure 2A.

**Figure 3.**
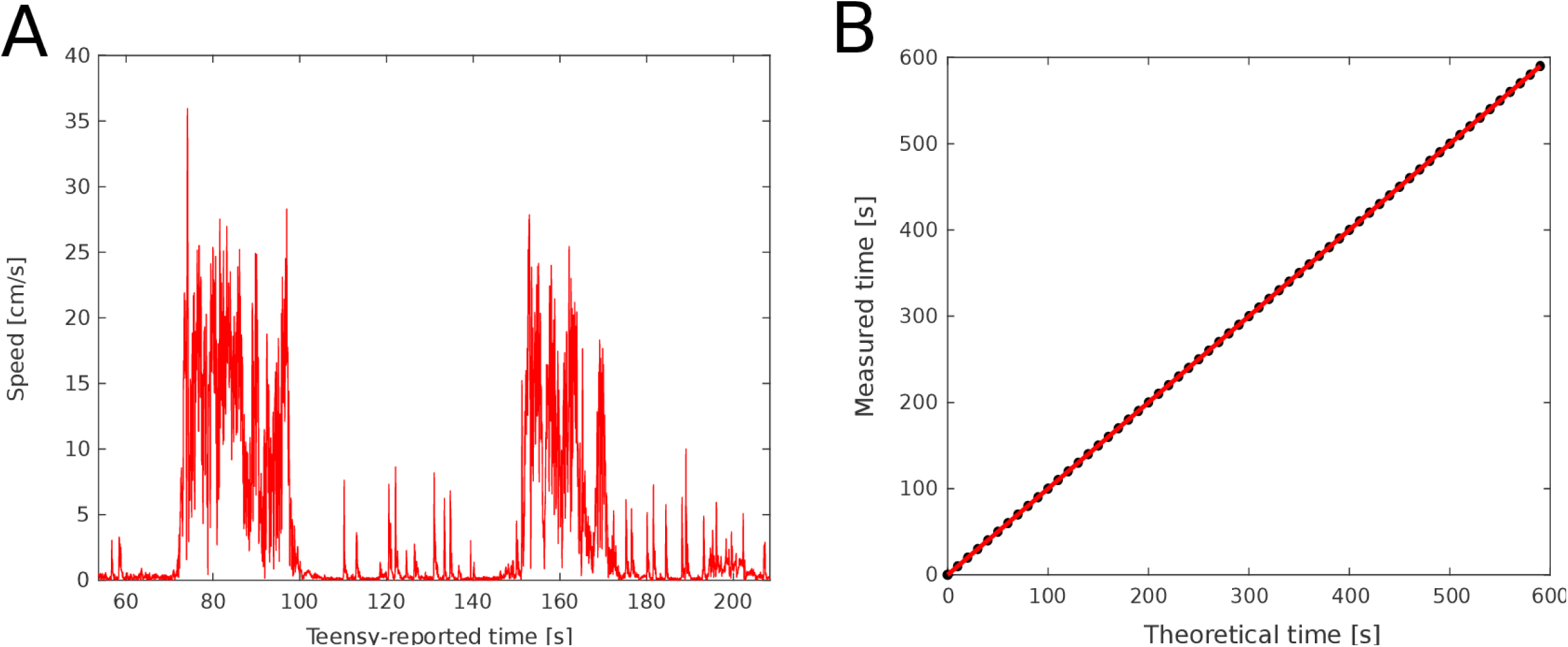
Temporal precision of the digital outputs in the motion tracking experiment. **A** Example recording of a head-fixed mouse running on the spherical treadmill. **B** Timing of digital pulses generated by the Teensy interface vs theoretical times of the digital pulses at exactly 20 Hz. Red indicates linear model prediction, and black are experimental data downsampled by a factor of 200 for visualization. The linear model estimates a slope of 1.000028937 ± 0.000000002 (t(11998) = 4.9e+08, p < 0.001, R^2^=1; intercept = 0.0007593 ± 0.0000007, t(11998) = 1.1e+03, p < 0.001).

We calculated the velocity of the mouse, which averaged 2.16 ± 4.46 cm/s over the 10 minute period (mean ± std, n=12000 time points) with a maximum velocity of 35.9 cm/s, in general agreement with velocities reported for head-fixed mice running on a spherical treadmill (Dombeck, et al. 2007)).

To characterize the temporal precision of the Teensy interface, we measured the timing of the Teensy digital output, and compared it to the theoretical 20 Hz signal using a linear model. We found that digital outputs have a near-perfect linear relationship with the theoretical signal (Figure 3B). However, we noted a 28.9 µs per second positive drift, resulting in an actual frequency of 19.999 Hz instead of 20.000 Hz. To further examine whether this small timing drift depends upon the frequency of data acquisition or the timing of the digital outputs, we performed 5 minute long recording sessions without a live mouse at 20, 50, and 100 Hz, and a 0.5ms long digital pulse designed to trigger image capture from the camera. We found that the actual frequencies were 19.999, 49.999, and 99.997 Hz, respectively. These all correspond to an approximately 30 µs delay per second, suggesting that the timing drift is independent of the data acquisition rate and may reflect the processor timing of the Teensy microcontroller. However, because motion sensor data are monitored with respect to the Teensy’s timing, the animal’s locomotion data readings remain precisely aligned to the time when image frame capture occurs.

Having assessed the timing of the digital output, we next quantified its temporal variation. We calculated the root mean squared error (RMSE) of the difference between the recorded timing of each digital pulse and the times predicted from the linear model. The RMSE was 42.7 µs, computed manually. Together, these results demonstrate that the Teensy interface timed by the “IntervalTimer” function can be used to generate digital pulses for pre cise image frame capture during behavioral experiments, while maintaining alignment of imaging data with behavioral parameters.

### 3.2 Trace eye blink conditioning behavioral experiment

In a second experiment, we reconfigured the Teensy interface for a trace conditioning eye blink learning experiment (Figure 1B and 2B), where a mouse can be trained to associate a conditioned stimulus (700ms long tone) with a subsequent unconditioned stimulus (a 100ms long gentle eye puff), separated by a brief memory trace time window (250ms). This experiment consisted of 50 trials, each lasting 20 seconds. We first characterize the temporal precision of the Teensy interface in a manner similar to that described in the motion tracking experiment. We recorded the timings of the digital pulses generated to trigger each image frame capture (Figure 4A), and detected a 33.4 microsecond delay per second. Thus, in this experiment, the Teensy interface has an actual frequency of 19.999 Hz instead of 20.000 Hz, identical to that observed in the motion tracking experiment. The RMSE of the Teensy interface is 13.3 µs computed manually, also similar to that observed in the motion tracking experiment.

**Figure 4.**
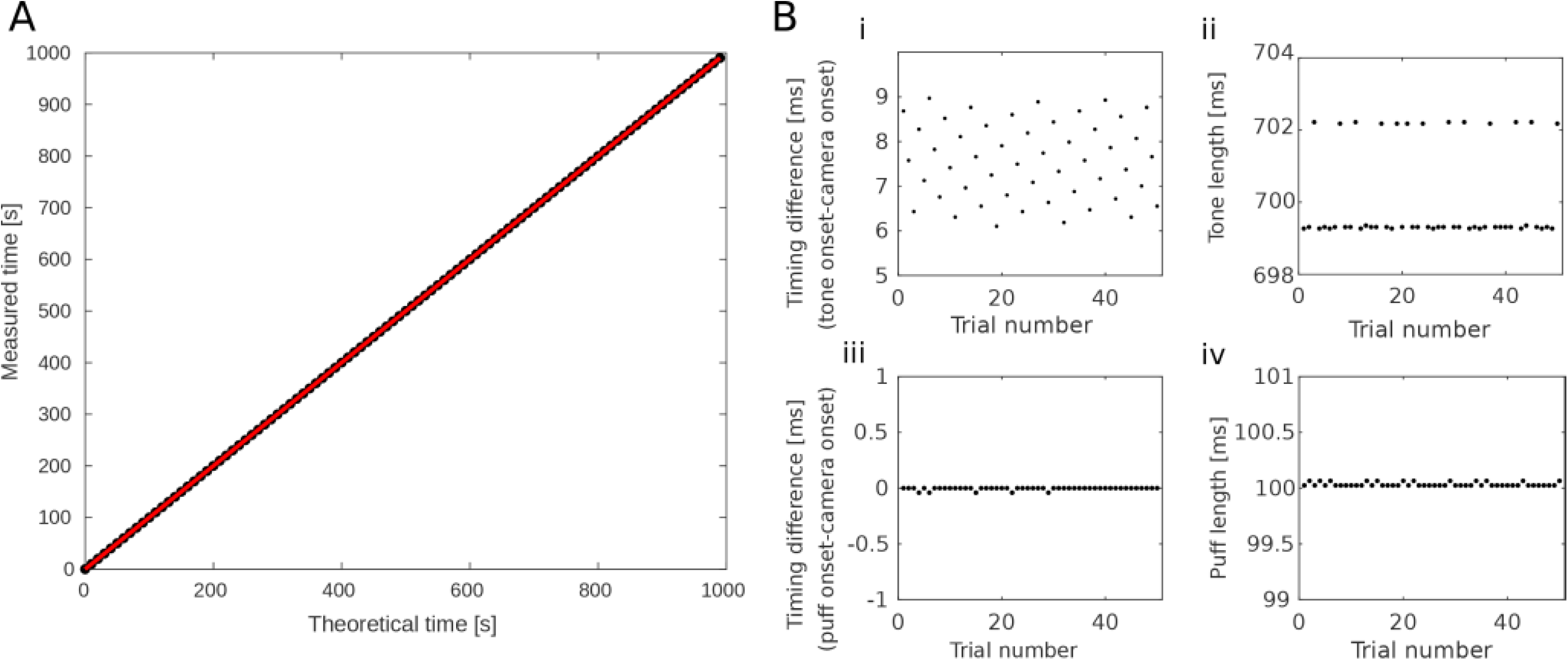
Temporal precision of the digital and analog outputs in the trace conditioning eye blink experiment. **A** Timing of the digital pulses generated by the Teensy interface vs theoretical times of the digital pulses at exactly 20 Hz. Linear model fit is shown in red, and in black are experimental data down-sampled by a factor of 200 for visualization. (R^2^=1, slope: 1.0000334 ± 0 (to machine precision), t(19998)=infinite, p<0.001). **B.** Timing of the analog output directed to the prop shield to generate an amplified auditory stimulus (i-ii) and the digital output directed to device to generate eye puff (iii-iv), both measured over 50 trials. (i) the difference between the onset of the analog output and the onset of the corresponding camera-directed digital pulse (mean=7.6 ± 0.9 ms, range=2.9 ms); (ii) the duration of the auditory stimulus across all trials (mean=700 ± 1 ms, range=2.9 ms, n=50 trials); (iii) the difference between the puff digital pulse and the camera-directed digital pulse, (mean = −0.004 ± 0.012 ms, range=0.04 ms); (iv) the duration of the puff digital pulse (100.03± 0.02 ms, mean ± std, n=50 trials).

We then characterized the precision of multiple digital outputs, by calculating the time difference between the digital pulses generated to drive eye puff versus the sCMOS camera (Figure 4Bii). We found that there was nearly no temporal difference between the onset of these two digital outputs (−0.004 ±0.012 ms,mean ± std, n=50 digital pulses). Similarly, the duration of the puff digital pulse was within 0.03 ms of the commanded duration of 100ms (100.03± 0.02 ms (mean ± std, n=50 digital pulses).

We next characterized the temporal precision of the analog output generated by the Teensy. We measured the analog output of the Teensy with the commercial TDT RZ5D recording device sampled at 24414.0625 Hz. Since analog outputs were generated together with the onset of the digital outputs designed to trigger camera image frame capture, we calculated the time difference between the onset of the analog output and the onset of the digital pulse (Figure 4Bi, for details see Methods). We found that the analog output lagged the digital output by 7.6 ± 0.9 milliseconds (mean ± std, n=50 pulses, Figure 4Bi). This delay is comparable to that reported using a different configuration of the Teensy to play a sound (Solari, et al. 2018). The duration of the tone remained equal to 700 ± 1 ms, (mean +/- std, n=50 digital/analog pulses Figure 4Bii), equivalent to the commanded duration of 700ms. Together, these results demonstrate that the Teensy interface, timed by the “elapsedMicros” function, is capable of generating digital and analog output with microsecond temporal precision.

To further examine whether this delay was related to our implementation of the Audio library or from writing to the analog pin itself, we directly generated an analog pulse without the Audio library from the analog pin using the Arduino command “analogWrite(A14, 4050)”. “A14” corresponds to the analog pin, and 4050 is a relative voltage level large enough to be recorded as a pulse by the TDT RZ5D system. We initiated 50 trials consisting of 50 millisecond long pulses through a digital pin and through the analog pin. Pulses to these two pins were programmed to occur near-simultaneously. We found that the analog output lagged the digital output by 0.8 + 5.8 µs (mean ± std, n=50 trials), suggesting that writing to the analog pin cannot account for the auditory signal delay generated through the Audio library. Thus the delay is due to the specific implementation of the audio library, and future changes to the Audio library could improve the temporal precision.

## 4. Conclusion and Discussion

In both experiments, the Teensy interface generated precisely timed digital pulses that can be used to control individual frame capture from a sCMOS camera at 20Hz. We detected a small drift of approximately 30 µs per second, suggesting an actual frequency of 19.999 Hz instead of the commanded 20Hz. This small 0.003% drift of the Teensy processing clock is linear, and can thus be calibrated if desired. This finding underscores the importance of having a highly precise central timer in each experiment. Synchronizing different devices only at the start of an experiment can lead to undesired temporal drifts, particularly in long experiments. While MATLAB or other PC-based programs can be programmed to control experimental timing, they may introduce timing jitter due to the demands of many PC system operations. Such timing jitter may have a significant impact depending on the study, especially when milliseconds time scale resolution is desired in systems neuroscience experiments.

Temporal accuracy is often important for animal behavioral training. For example, a precisely timed conditioned stimulus (tone) and unconditioned stimulus (puff) are important for animals to build association in trace conditioning eye blink experiments. We demonstrate that the Teensy interface can accurately generate multiple digital pulses to drive different devices, including the tone, the puff and the sCMOS camera. Additionally, we demonstrate that the Teensy interface precisely delivered longer duration digital and analog pulses, such as the tone that lasted for 700ms in the trace conditioning eye blink experiment. These results demonstrate that Teensy interface is a viable, inexpensive alternative that is also able to simultaneously capture imaging data using our simple software functions.

A major advantage of the Teensy over Arduino Uno microcontrollers is its ability to generate a true, 12 bit analog signal. While Arduino Uno microcontrollers can generate an analog-like signal via pulse-width modulation, this output is a square wave. We used the Teensy interface to deliver an auditory stimulus through the built-in Audio library, and our analog output showed a 7.6ms delay. This small delay is due in large part to the implementation of the Audio library. It is possible that other ways of utilizing the analog output would allow the generation of more temporally precise audio signals. However, altering the amplitude of a single sine wave via the Audio library is easy to implement, utilizing only a few lines of code within a single script.

### 4.1 Conclusion

We demonstrate a Teensy 3.2 interface capable of integrating a sCMOS camera into two behavioral experimental settings. In one setting, the Teensy interface simultaneously generates digital pulses that can be directed for individual frame capture from a sCMOS camera, while simultaneously tracking an animal’s locomotion using recently developed high precision ADNS-9800 gaming sensors. The easy integration of the sCMOS camera and the ADNS-9800 sensors illustrates the flexibility of the Teensy interface in designing experiments that require novel instrumentation. In the second experiment, we demonstrate that the Teensy interface, in conjunction with a prop shield, is capable of generating both analog and digital outputs with precise timing during an eye blink trace conditioning experiment. We characterized two timer functions, “IntervalTimer” and “elapsedMicros”, both of which offered equivalent microsecond temporal precision, and “elapsedMicros” additionally allows access to the Audio library. Thus the Teensy interface, a Teensy 3.2 and custom software functions, provides a user-friendly, easily adaptable, and temporally precise platform for integrating sCMOS cameras into behavioral experimental designs. This Teensy interface can be immediately adopted for the motion tracking and the trace conditioning eye blink behavioral experiments demonstrated here, or can be customized for other types of behavioral experiments where sCMOS camera-based imaging is desired.

## Acknowledgements

M.F.R. performed data analysis. M.F.R. and H.J.G. conducted the motion tracking experiment. M.F.R. conducted the trace conditioning eye blink experiment. M.F.R., M.B., and D.R.M. wrote the software. M.F.R., M.B., D.R.M., and R.K. contributed to the Teensy interface conceptualization. M.F.R., H.J.G., and X.H. wrote the manuscript. X.H. supervised the study. The authors would also like to acknowledge Thomas Romano for helpful conversations, and users “Theremingenieur” and “PaulStoffregen” from the PJRC forums (https://forum.pjrc.com/) for responding to questions relating to the trace eye blink conditioning experiment.

The authors have no competing financial interests.

